# A Novel Optimized Silver Nitrate Staining Method for Visualizing the Osteocyte Lacuno-Canalicular System

**DOI:** 10.1101/2025.01.14.632997

**Authors:** Jinlian Wu, Chunchun Xue, Qiang Li, Hongjin Wu, Jie Zhang, Chenglong Wang, Weiwei Dai, Libo Wang

## Abstract

A 50% silver nitrate solution is commonly used to stain and characterize the osteocyte lacuno-canalicular system (LCS) in bone biology research. However, variability in reagent concentrations and types, along with inconsistent staining procedures, have limited the broader application of this method in osteocyte research. In this study, we present a novel optimized silver nitrate staining technique aimed at addressing these limitations. This new method utilizes a 1 mol/L silver nitrate solution in combination with a type-B gelatin-formic acid solution at various concentrations (0.05%-0.5% gelatin and 0.05%-5% formic acid, or 1%-2% gelatin and 0.1%-2% formic acid) in volume ratios of 4:1, 2:1, or 1:1, or a 0.5 mol/L silver nitrate solution at a 4:1 ratio. The staining process is carried out for 1 hour under ultraviolet light or 90 minutes under regular room light (or dark), followed by washing with Milli-Q water to terminate the reaction. We applied this new method to stain the osteocyte LCS in bone samples from different species and pathological bone models. The technique consistently produced clear, distinct staining patterns across all samples. Moreover, our novel method revealed a greater number of LCS compared to the traditional 50% silver nitrate solution. This suggests that the commonly used 50% silver nitrate method may disrupt or inadequately reveal the LCS in certain species, potentially leading to an underestimation of LCS density and number. In conclusion, our novel silver nitrate staining method provides a simpler and more cost-effective alternative to the traditional technique. By offering a more accurate and comprehensive analysis of the LCS across species, this approach has the potential to advance research on osteocyte morphogenesis, as well as the functional and evolutionary adaptations of the osteocyte LCS across different taxa.

## Introduction

Osteocytes are terminally differentiated osteoblasts embedded in the bone matrix within small cavities called lacunae. A hallmark of osteocytes is their dendritic processes—long, branching protrusions that extend from the cell body. These dendrites pass through tiny canals called canaliculi, which connect the lacunae throughout the bone matrix [1–3]. This network of lacunae and canaliculi, known as the lacuno-canalicular system (LCS), along with the dendritic network, exhibits a remarkable and intricate architecture similar to the dendritic connectivity observed in neuronal cells [4–5]. The adult human skeleton contains approximately 42 billion osteocytes and 3.7 trillion dendrites, with each osteocyte having 18 to 106 dendrites that extend into 53 to 126 canaliculi connected to the lacunae [4].

The osteocytes and their LCS are critical for several physiological functions, including nutrient and waste exchange, as well as cell signaling between osteocytes and other cells, such as osteoblasts, osteoclasts, and vascular endothelial cells [6–9]. Given the central role of the LCS in bone biology, extensive research has been dedicated to understanding its structural and functional changes in various contexts, including bone development, bone diseases (e.g., osteoporosis, osteoarthritis), pregnancy, and aging [10–14].

Among the various histological staining techniques, the silver nitrate method, particularly the Ploton method developed in 1986, is widely used to investigate the osteocyte LCS [15–17]. This technique is effective in revealing the structure of the LCS, allowing for detailed analysis of osteocyte connectivity. However, despite its utility, the method has several limitations. These include inconsistencies in the concentrations of silver nitrate and gelatin-formic acid solutions, a lack of standardization in staining protocols, variability in the type of gelatin used, and inconsistencies in staining and termination conditions. These factors have impeded the wider adoption of Ploton method in osteocyte research.

In this study, we developed a simplified, more environmentally friendly silver nitrate staining method that improves upon the classical method that uses a 50% silver nitrate solution. Our approach used a 1 mol/L silver nitrate solution combined with type-B gelatin-formic acid solutions at various concentrations (0.05%-0.5% gelatin and 0.05%-5% formic acid, or 1%-2% gelatin and 0.1%-2% formic acid) in volume ratios of 4:1, 2:1, or 1:1, or a 0.5 mol/L silver nitrate solution at a 4:1 ratio. The staining was performed for 1 hour under UV light or 90 minutes under regular room light (or dark). After staining, the slides were washed with Milli-Q water (no need for 5% Sodium thiosulfate), followed by standard dehydration, clearing, and mounting procedures. The novel optimized method yielded consistent and distinct staining across different species and disease models, demonstrating superior efficiency in staining osteocyte LCS at higher densities and greater numbers compared to the Ploton method. We believe that our new method will contribute to advancing the study of bone biology and have broader applications in the diagnosis and treatment of bone-related diseases.

## Materials and Methods

### Animal bone sample collection

The Male C57BL/6J mice (4-8 weeks old) and Sprague-Dawley rat (8 weeks old) were purchased from Hangzhou Ziyuan Laboratory Animal Technology Co., Ltd (Hangzhou, China). Prrx1Cre mice and Zeb1^fx/fx^ mice were purchased from GemPharmatech Co., Ltd (Jiangsu, China). All mice and rat were housed under standard conditions at the animal facility of Shanghai Municipal Hospital of Traditional Chinese Medicine. Rabbit (*Oryctolagus cuniculus*), common carp (*Cyprinus carpio*), silver carp (*Hypophthalmichthys molitrix*), snakehead (*Channa maculata*), little yellow croaker (*Larimichthys polyactis*), bullfrogs (*Lithobates catesbeiana*), chickens (*Gallus gallus domesticus*), and wall lizards (*Gekko sp.*) were purchased from a local food market and an online traditional Chinese medicine herb store. A bone growth retardation model was created by administering dexamethasone (2 mg/kg/day) intraperitoneally to 4-week-old male mice for four weeks. A transgenic mouse model (Prrx1Cre;Zeb1*^fx/fx^*), which exhibits pathological features of osteogenesis imperfecta and specifically overexpresses Zeb1 in skeletal mesenchymal stem cells, was generated by crossing Prrx1Cre mice with Zeb1^fx/fx^ mice. Femurs from mice and rats were collected after euthanasia via carbon dioxide inhalation. Femurs from bullfrogs, chickens, rabbits and wall lizards, as well as caudal vertebrae from common carp, silver carp, snakehead, and little yellow croaker, were isolated upon purchase. Animal experiments were performed in strict compliance with the guidelines and were approved by the Animal Experiments Ethical Committee of Shanghai Municipal Hospital of Traditional Chinese Medicine (Approval No. 2020007).

### Reagents and Apparatus

Reagents: 10% EDTA Decalcification solution(E671001-0500, Sangon Biotech), 10% Neutral formalin fixative(311010014, Wexis), 1 mol/L Silver nitrate solution(P1929026, Bolinda Technology), Silver nitrate power(C510027-0010, Sangon Biotech), Type-A Galetin(A609764-0100, Bloom 238∼282, Sangon Biotech), Type-B Galetin(G8061, bloom 225, Solarbio), 88% Formic acid(10010118, SinoPharm), Dexamethasone(HY-14648, MedChemExpress), Sodium thiosulfate(A600484-0500, Sangon Biotech), Neutral balsam mounting medium(G8590, Solarbio). Apparatus: Biological Safety Cabinet(1379/1389 with 254nm UV light G3675L 31∼40W, Thermo), Automatic Benchtop Tissue Processor(TP1020, Leica), Tissue Embedding Station(EG1160, Leica), Rotary microtome(RM2135, Leica), Histology Water Bath(Histobath HI1210, Leica), Ultrapure Water Purification System(Milli-Q Elix® Essential, Millipore), Nikon bright-field microscope(ECLIPSE 200, Nikon).

### Bone sample processing

All animal bone samples were fixed in 10% neutral formalin for 24 hours, then washed in 18.2 MΩ·cm Milli-Q water for 60 minutes. The bones were subsequently decalcified in 10% EDTA (pH 7.4) at room temperature for 28 days, with the solution replaced every three days. After decalcification, the bones were dehydrated, paraffin-embedded, sectioned at 4 μm thickness, and stored at 4°C.

### Silver nitrate staining

Solution preparation: (1) 1 mol/L Silver nitrate solution: Prepare the silver nitrate solution using 18.2 MΩ·cm Milli-Q water. Store the solution at room temperature, protected from light. (2) Gelatin-Formic acid solution: Prepare a 1% (v/v) formic acid solution using 18.2 MΩ·cm Milli-Q water. Weigh 2% (w/v) type-B gelatin and add it to the formic acid solution. Heat the mixture at 37°C for 2 hours, or until the gelatin is fully dissolved. Store at room temperature. (3) Staining solution: Combine two volumes of the 1 M silver nitrate solution with one volume of the 2% type-B gelatin-1% formic acid solution. Prepare the staining solution immediately before use.

Staining procedure: Following deparaffinization with xylene and rehydration through a graded alcohol series, paraffin sections are washed twice in 18.2 MΩ·cm Milli-Q water for 5 minutes each. Next, two volumes of a 1 mol/L silver nitrate solution are mixed with one volume of a 2% (w/v) type-B gelatin-1% (v/v) formic acid solution and applied to the slides. The slides are then stained for 60 minutes under 254 nm ultraviolet light or 90 minutes under regular room light at room temperature. After staining, the reaction is terminated by washing the slides twice with 18.2 MΩ·cm Milli-Q water for 5 minutes each, followed by dehydration, clarification, and mounting with neutral balsam. The general procedure for the optimized silver nitrate staining method for osteocyte LCS is shown in Figure 7.

### LCS Quantification

The osteocyte LCS area was quantified and normalized to the total bone area as previously described [18–19]. Images at 400× magnification were acquired using a Nikon ECLIPSE 200 bright-field microscope (Nikon, Japan) and analyzed with ImageJ software (NIH, USA) by thresholding grayscale images.

### Statistics analysis

Data were presented as means ± standard deviation, shown by error bars. Comparisons were performed using either one-way ANOVA or an unpaired two-tailed Student’s t-test in GraphPad Prism 8.0.1 software. A p-value of < 0.05 was considered statistically significant.

## Results

### 1. A 1 mol/L or 0.5 mol/L silver nitrate solution is an effective stain for osteocyte LCS

In preliminary experiments, we unexpectedly found that a 1 mol/L silver nitrate solution could effectively stain the osteocyte lacunae-canalicular system (LCS). Building on this discovery, we systematically evaluated the effects of varying silver nitrate concentrations (3 mol/L, 2 mol/L, 1 mol/L, 0.5 mol/L, 0.25 mol/L, and 0.1 mol/L) combined with a 2% type B gelatin and 1% formic acid solution at different volume ratios (4:1, 2:1, 1:1, 1:2, and 1:4). Staining was performed under UV light for either 1 or 2 hours, followed by termination with Milli-Q water (Figure 1).

**Figure 1.**
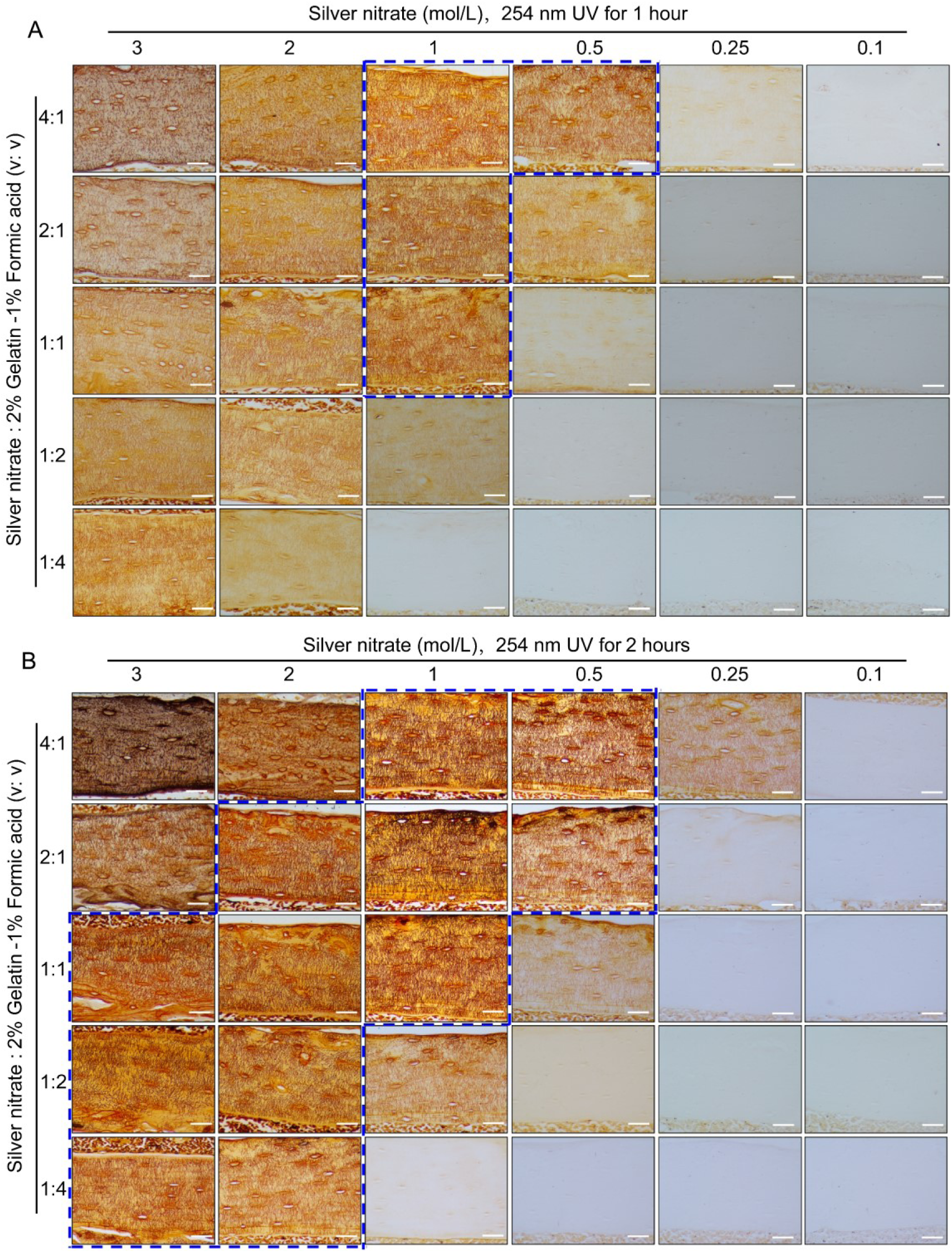
Staining effects of different concentrations of silver nitrate combined with different volume ratios with 2% gelatin-1% formic acid solution when staining under UV 1hour or 2 hours on LCS. (A) Effects of exposure to 254 nm UV light for 1 hour to stain LCS. Scale bars = 50 μm. (B) Effects of exposure to 254 nm UV light for 2 hours to stain LCS. Scale bars = 50 μm. The dashed blue box indicates the optimal silver nitrate concentration and the volume ratio of silver nitrate solution to type-B gelatin-formic acid solution for clear and effective staining of osteocyte LCS. All scale bars = 50 μm.

Staining results indicated that under 254 nm UV light for 1 hour (Figure 1A), the 1 mol/L silver nitrate solution with 2% gelatin and 1% formic acid at ratios of 4:1, 2:1, and 1:1, as well as the 0.5 mol/L silver nitrate solution at a 4:1 ratio, produced clear and effective LCS staining compared to other combinations. When stained for 2 hours under 254 nm UV light (Figure 1B), the effective staining range expanded to include: 3 mol/L silver nitrate with 2% gelatin and 1% formic acid at ratios of 1:1, 1:2, or 1:4; 2 mol/L silver nitrate with 2% gelatin and 1% formic acid at ratios of 2:1, 1:1, 1:2, or 1:4; 1 mol/L silver nitrate at ratios of 4:1, 2:1, or 1:1; and 0.5 mol/L silver nitrate at ratios of 4:1 and 2:1.

Regardless of the staining duration (1 or 2 hours), the use of 3 mol/L silver nitrate with 2% gelatin and 1% formic acid at ratios of 4:1 and 2:1, or 2 mol/L silver nitrate at a 4:1 ratio, resulted in unclear staining and a darker background. This led to intermittent staining patterns, with fewer LCS stained compared to those stained with the 1 mol/L silver nitrate solution. Additionally, silver nitrate concentrations of 0.25 mol/L and 0.1 mol/L were ineffective in staining the osteocyte LCS under the same conditions. These findings suggest an optimal range of silver nitrate concentration for effective LCS staining; both excessively high and low concentrations failed to produce clear staining and accurate quantification of the LCS.

Based on the qualitative assessment of LCS staining, our data indicate that a 1 mol/L silver nitrate solution combined with 2% gelatin and 1% formic acid at ratios of 4:1, 2:1, or 1:1, or a 0.5 mol/L silver nitrate solution at a 4:1 ratio, under 254 nm UV light for 1 hour, can clearly and effectively stain the osteocyte LCS (as indicated by the dashed blue box in Figure 1).

### 2. A 1 mol/L silver nitrate solution, when combined with gelatin concentrations ranging from 0.05% to 0.5% (w/v) and formic acid concentrations from 0.05% to 5% (v/v), or with 1%-2% (w/v) gelatin and 0.1% to 2% (v/v) formic acid, effectively stains osteocyte LCS

After establishing the effective range of silver nitrate, we evaluated the impact of varying concentrations of type-B gelatin (0%-5%, w/v) and formic acid (0%-10%, v/v) on LCS staining, using a 1 mol/L silver nitrate solution and a gelatin-formic acid solution at a 2:1 volume ratio. The staining was conducted under 254 nm UV light for 1 hour, followed by termination with Milli-Q water (Figure 2).

**Figure 2.**
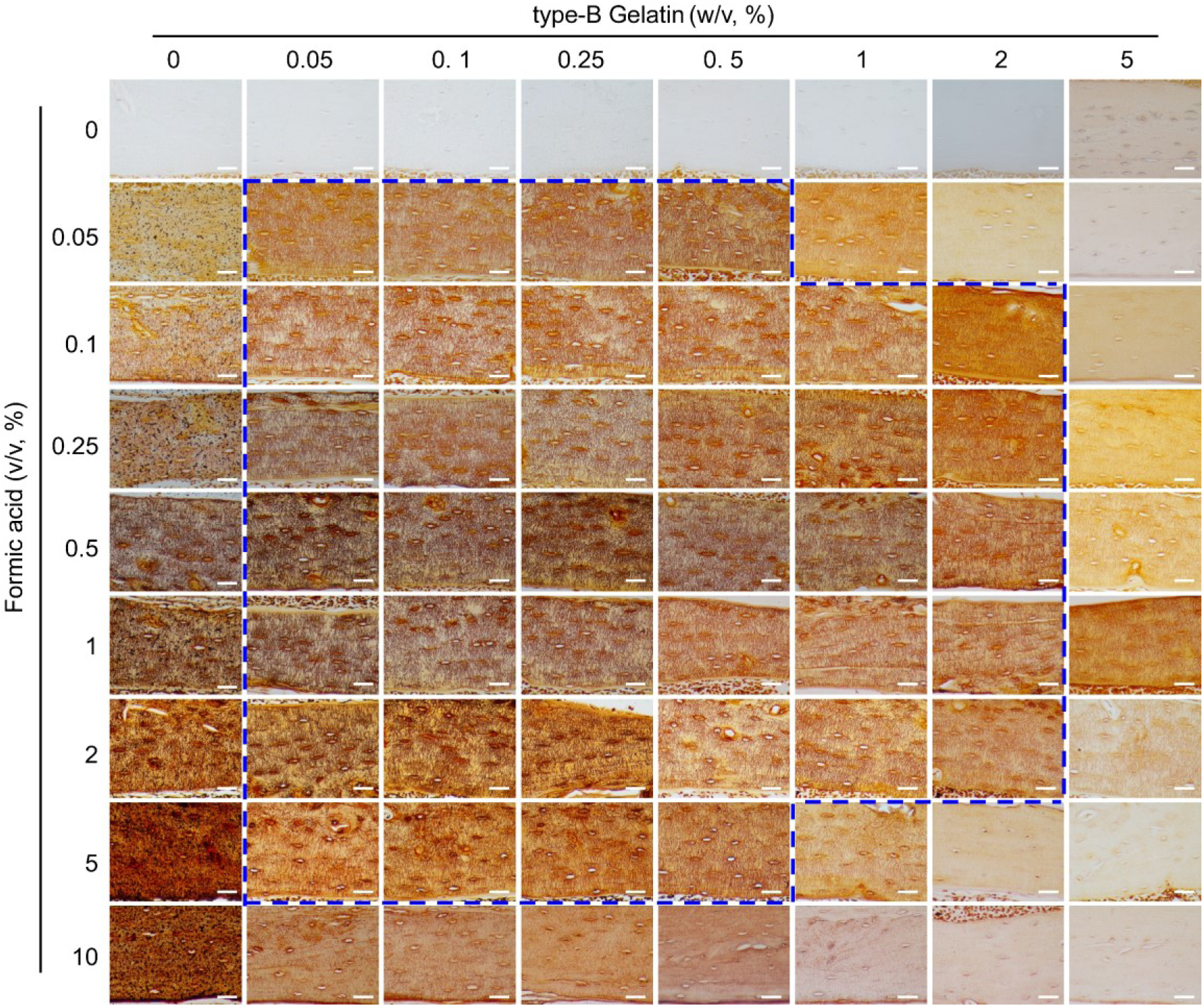
Comparison of the staining effects on LCS using gelatin (0%-5%, w/v)-formic acid (0%-10%, v/v) solutions at different concentrations, in a 2:1 volume ratio with 1 mol/L silver nitrate. The dashed blue box indicates the effective concentration range of the type-B gelatin and formic acid solution combination. All scale bars = 50 μm.

The results indicated that silver nitrate alone could not stain LCS when both gelatin and formic acid concentrations were zero. When the gelatin concentration was zero and formic acid was ≥ 0.05%, LCS structures were visible under the microscope after staining, but a significant number of black granular precipitates adhered to the tissue, rendering the data unusable. In the absence of formic acid, silver nitrate failed to stain LCS at gelatin concentrations of 0.05% or higher. Furthermore, at a specific formic acid concentration (≥ 0.05%), the intensity of LCS staining decreased as gelatin concentration increased. Conversely, at a specific gelatin concentration (≥ 0.05%), the intensity of LCS staining initially increased and then decreased with increasing formic acid concentration.

In summary, the findings demonstrated that solution ratios with gelatin concentrations ranging from 0.05% to 0.5% and formic acid concentrations from 0.05% to 5%, as well as 1%-2% gelatin and 0.1% to 2% formic acid, resulted in the clear LCS staining, as indicated by the dashed blue box in Figure 2.

### 3. Optimal staining of the LCS can be achieved using type-B gelatin, with staining conducted under UV light for 1 hour or in room light (or in the dark) for 90 minutes, followed by termination of the reaction with Milli-Q water

Notable differences exist in the silver nitrate staining methods reported across various studies, particularly regarding the type of gelatin used (type A [19] or type B [20]), lighting conditions (UV, regular room light, or dark [17]), and reaction termination methods (Milli-Q water [15] or 5% sodium thiosulfate [16]). To evaluate the effects of these factors on LCS staining, we employed a 1 mol/L silver nitrate solution with 2% gelatin and 1% formic acid in a 2:1 volume ratio, staining under UV light for 1 hour (Figure 3).

**Figure 3.**
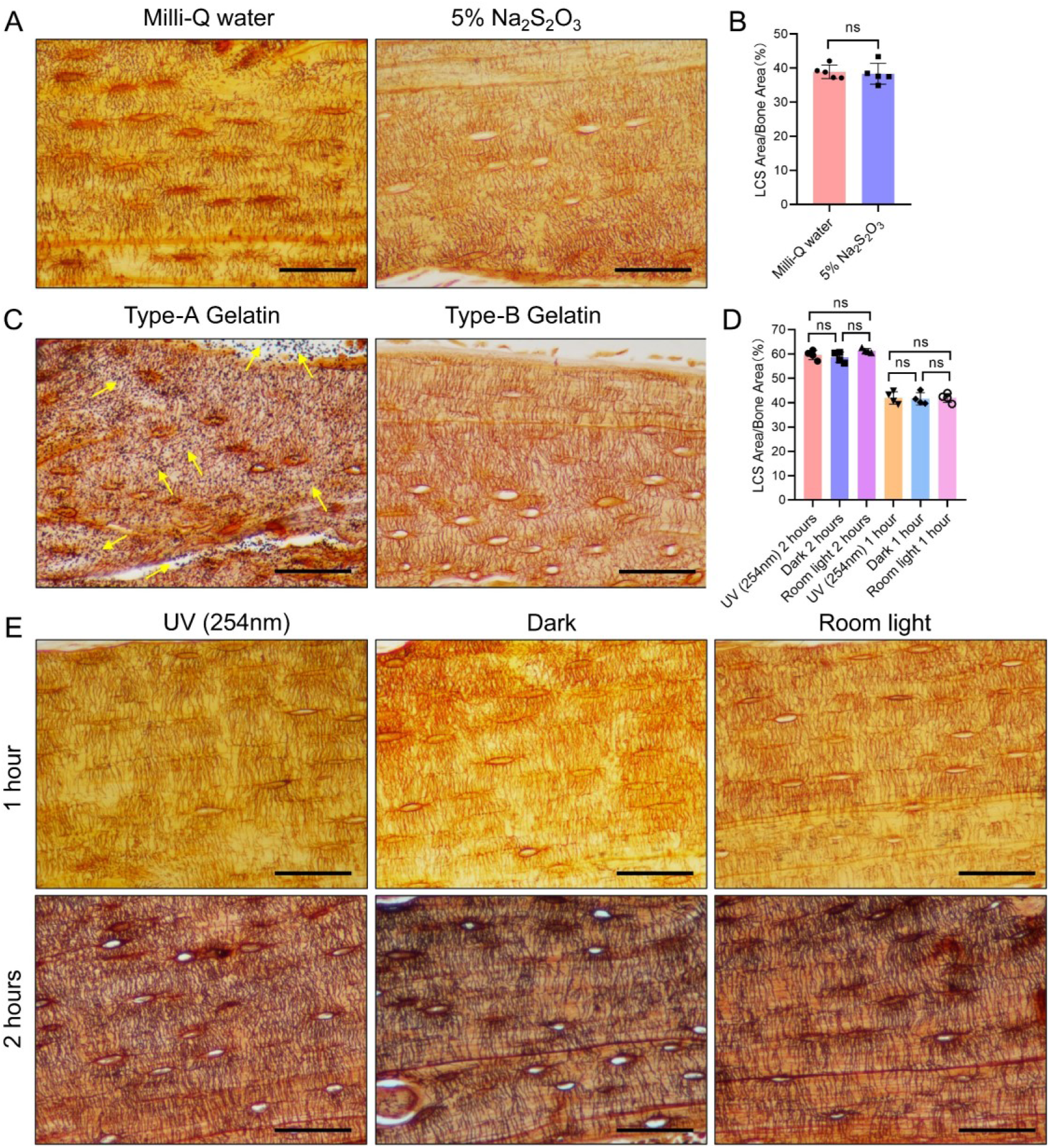
Effects of Three Different Conditions on Silver Nitrate Staining of LCS. (A) Effects of post-reaction Mill-Q water washing and 5% sodium thiosulfate washing on LCS staining. Scale bars = 50 μm. (B) Comparison of staining effects on osteocyte LCS between Milli-Q water and 5% sodium thiosulfate, analyzed using an unpaired two-tailed Student’s t-test. Data are presented as mean ± SD. ns p > 0.05, n = 5 for repeated staining with 3-4 regions of interest (ROIs) per staining section. (C) Effects of type-A and type-B gelatin on LCS staining. Yellow arrows indicate the presence of a large amount of black granular precipitates on the tissues after staining with type-A gelatin. Scale bars = 50 μm. (D) Comparison of staining effects on osteocyte LCS under three different lighting conditions, analyzed using one-way ANOVA. Data are presented as mean ± SD. ns p > 0.05, n = 4 for repeated staining with 3-4 ROIs per staining section. (E) Effect of different light conditions (UV, room light, or dark) on LCS staining. Scale bars = 50 μm.

The results indicated that tissues terminated with Milli-Q water exhibited clearer contrast and brighter staining compared to those terminated with 5% sodium thiosulfate (Figure 3A). However, no significant difference in the density of LCS staining was observed between the two termination methods (Figure 3B). When different types of gelatin were tested, type-A gelatin resulted in heavy black particulate deposits (Figure 3C, indicated by yellow arrows), while type-B gelatin produced a clear background without such deposits (Figure 3C). This suggests that the black particulate deposits observed in some studies may be due to the improper selection of gelatin type [19, 21]. Staining under different light conditions for one hour revealed that all three conditions (UV, regular room light, and dark) resulted in clearer LCS staining (Figure 3E). While no significant difference in LCS staining density was observed between the UV condition and regular room light (or dark) after one hour (Figure 3D), the UV condition produced deeper staining of the LCS compared to regular room light or dark under the same duration. After two hours of staining, the staining depth under regular room light and dark was comparable to that achieved with UV light (Figure 3E). Despite no noticeable difference in LCS staining density across the three light conditions after two hours (Figure 3D), the LCS background was slightly darker, and the contrast was less clear compared to one hour of staining.

Based on these observations, we conclude that staining for 60 minutes under UV light or for 90 minutes under regular room light or in the dark yields the most distinct staining. Therefore, optimal LCS staining on paraffin sections can be achieved using type-B gelatin, with staining for 1 hour under UV light or 90 minutes under regular room light or in the dark, followed by termination of the reaction with Milli-Q water.

### 4. A 1 mol/L silver nitrate solution effectively stains the LCS across different taxa

To assess the generalizability of the optimized staining method, we performed silver nitrate staining of LCS on skeletal samples from various species. The staining solution was prepared by mixing 1 mol/L silver nitrate with 2% type-B gelatin and 1% formic acid in a 2:1 volume ratio. The samples were reacted under UV light for 1 hour, followed by termination with a Milli-Q water wash (Figure 4).

**Figure 4.**
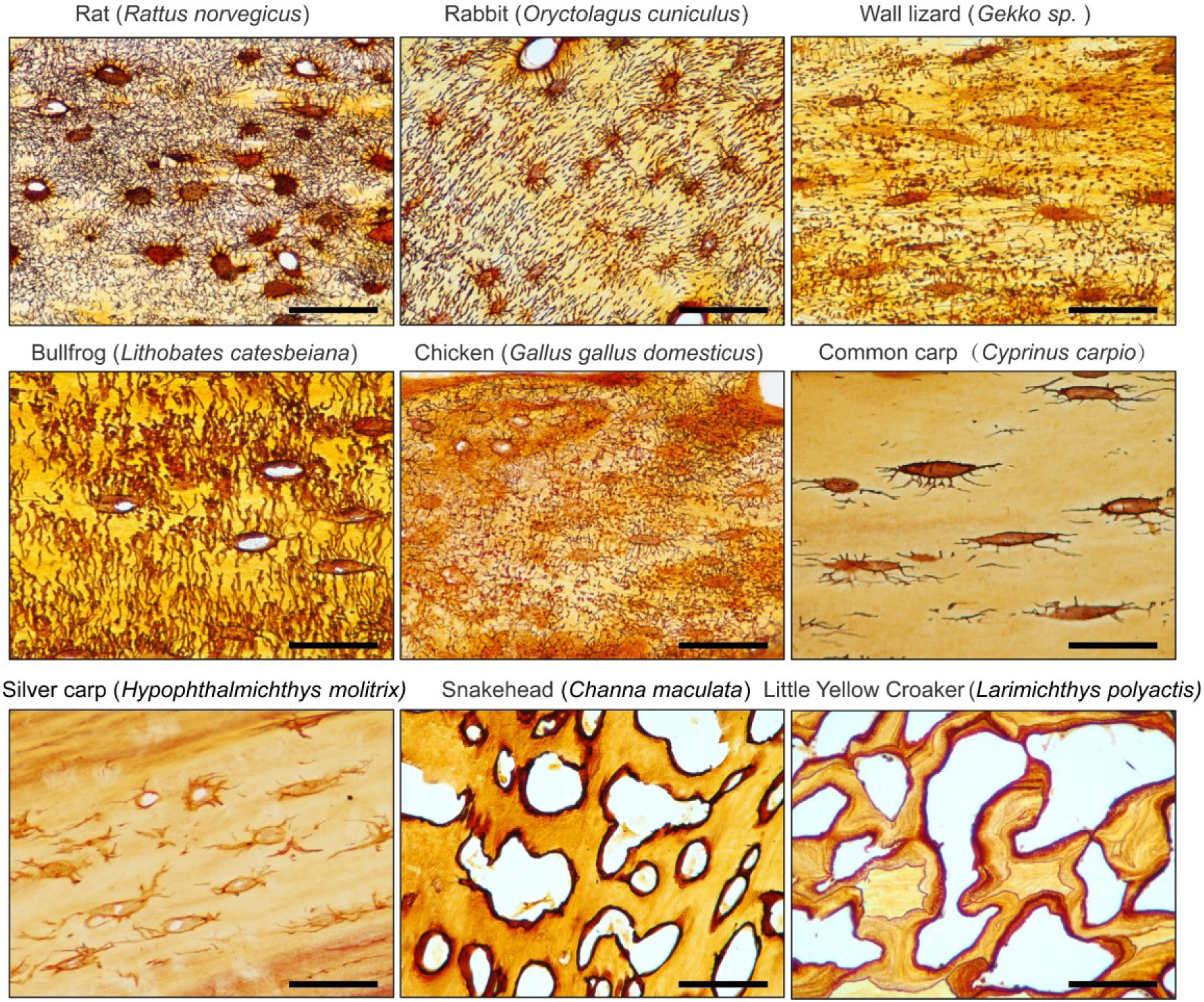
1 mol/L Silver Nitrate solution effectively stains the osteocyte LCS across different animal species. All Scale bars = 50 μm.

The results showed that the optimized method effectively stained LCS in mammals (e.g., rat, rabbit), amphibians (e.g., bullfrog), birds (e.g., chicken), reptiles (e.g., wall lizard), and fishes (e.g., common carp, silver carp, snakehead, and little yellow croaker). However, the number of osteocytes in the caudal vertebrae of teleosts, such as common carp (*Cyprinus carpio*) and silver carp (*Hypophthalmichthys molitrix*), was notably low, and the LCS was sparsely present. In contrast, in the caudal vertebrae of other teleosts, such as snakehead (*Channa maculata*) and little yellow croaker (*Larimichthys polyactis*), silver nitrate staining revealed that mature vertebrae in these species were completely devoid of osteocytes and canalicular structures, with only lacunae-like structures visible. This suggests that teleosts generally lack mature osteocyte dendritic structures and a well-developed LCS, likely as an adaptation to their buoyant aquatic environments, as shown by phylogenetic studies of acellular bone [22, 23].

Thus, our novel silver nitrate method demonstrated broad applicability to skeletal tissues from various animal groups, highlighting its potential for comparative evolutionary biology studies on osteocytes and LCS systems across species.

### 5. A 1 mol/L silver nitrate solution effectively stains osteocyte LCS in both pathological and transgenic mouse models

To assess the potential utility of the optimized method in pathological models, we performed silver nitrate staining on bone samples from two mouse models: a glucocorticoid-induced bone growth retardation model and a transgenic mouse model (Prrx1Cre; Zeb1^fx/fx^) that overexpresses the Zeb1 transcription factor specifically in bone mesenchymal stem cells, exhibiting an osteogenesis imperfecta phenotype. The staining solution was prepared by mixing 1 mol/L silver nitrate with 2% type-B gelatin and 1% formic acid in a 2:1 volume ratio, and the reaction was under UV light for 1 hour and terminated by Milli-Q water washing (Figure 5).

**Figure 5.**
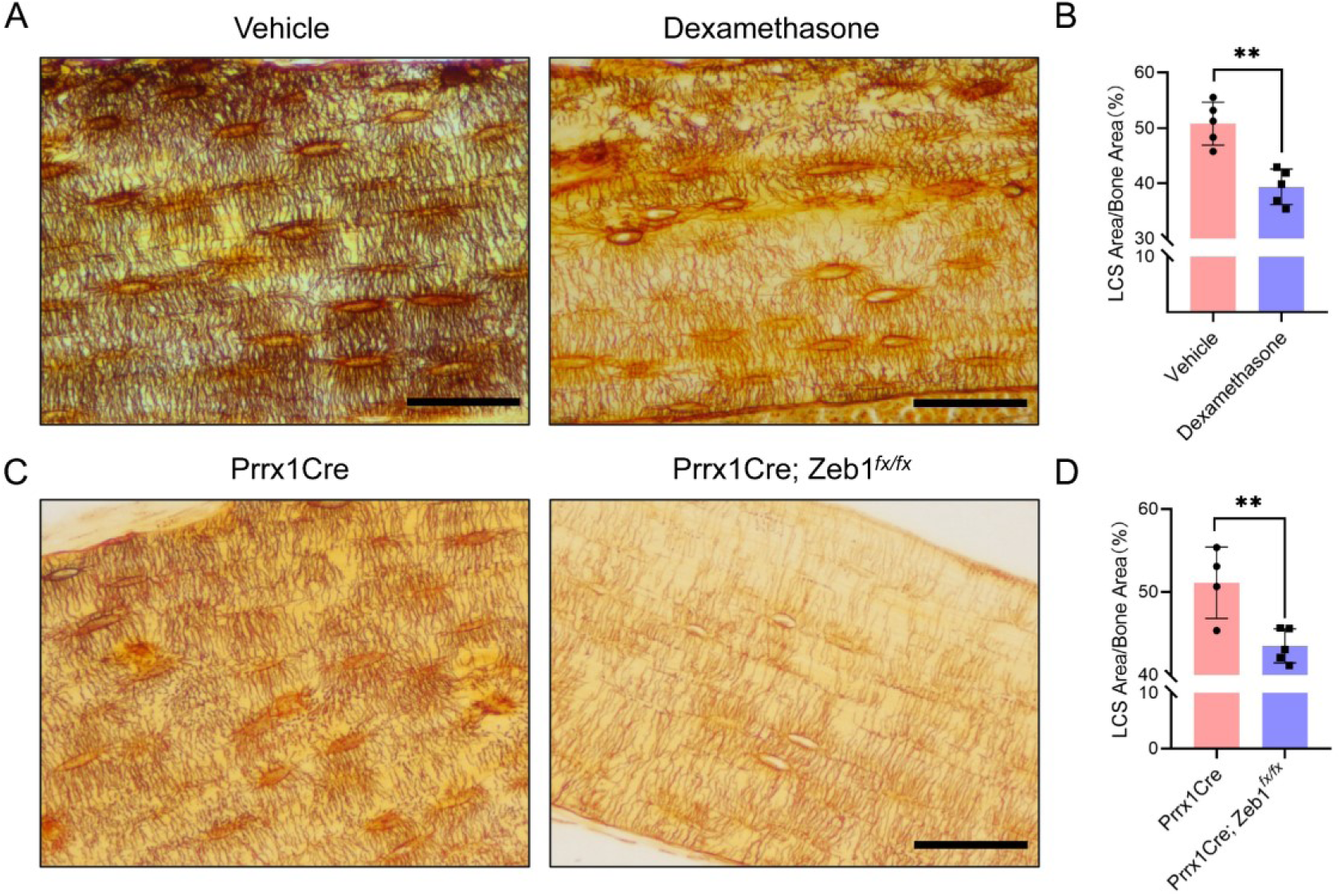
LCS Staining of Cortical Bone in Two Mouse Models Using 1 mol/L Silver Nitrate. (A) Cortical bone LCS was significantly reduced in glucocorticoid-induced bone growth retardation mice compared to vehicle-treated mice. Scale bars = 50 μm. (B) Comparison of osteocyte LCS density between glucocorticoid-induced bone growth retardation mice and vehicle mice, analyzed using an unpaired two-tailed Student’s t-test. Data presented as mean ± SD. **p < 0.01, n = 5 for repeated staining with 3-4 ROIs per bone section. (C) Cortical bone LCS was significantly decreased in Prrx1Cre;Zeb1*^fx/fx^* mice compared to Prrx1Cre mice. Scale bars = 50 μm. (D) Comparison of osteocyte LCS density between Prrx1Cre;Zeb1^fx/fx^ mice and Prrx1Cre mice, analyzed using an unpaired two-tailed Student’s t-test. Data are presented as mean ± SD. **p < 0.01, n = 4-5 for repeated staining with 3-4 ROIs per staining section.

The staining results showed that the optimized method effectively stained LCS in both models (Figure 5). The density of LCS was significantly reduced in the glucocorticoid-treated mice (Figure 5A and 5B) and Prrx1Cre; Zeb1^fx/fx^ mice (Figure 5C and 5D) compared to control animals. Notably, the staining results from Prrx1Cre; Zeb1^fx/fx^ mice indicated that Zeb1 overexpression might impair the differentiation of mesenchymal stem cells into osteoblasts and osteocytes, disrupting the balance of chondrocyte and osteoblast differentiation. This led to pathological cartilage remnants in the cortical bone of adult mice (Safranin O and Fast Green staining, data not shown). These findings further suggest that our optimized staining method can effectively identify changes in LCS in common or new animal pathology models.

### 6. A 1 mol/L silver nitrate solution stains osteocyte LCS more effectively than the Ploton method

Next, we compared the optimized method using a 1 mol/L silver nitrate with the classical Ploton silver nitrate staining method, which uses 50% silver nitrate (2.943 mol/L), for staining LCS. We tested different concentrations of silver nitrate solutions (1 mol/L and 50%), combined with a 2% Type-B gelatin and 1% formic acid solution at varying ratios (4:1, 2:1, and 1:1). The reactions were carried out under UV light for 1 hour, followed by washing with Milli-Q water (Figure 6).

**Figure 6.**
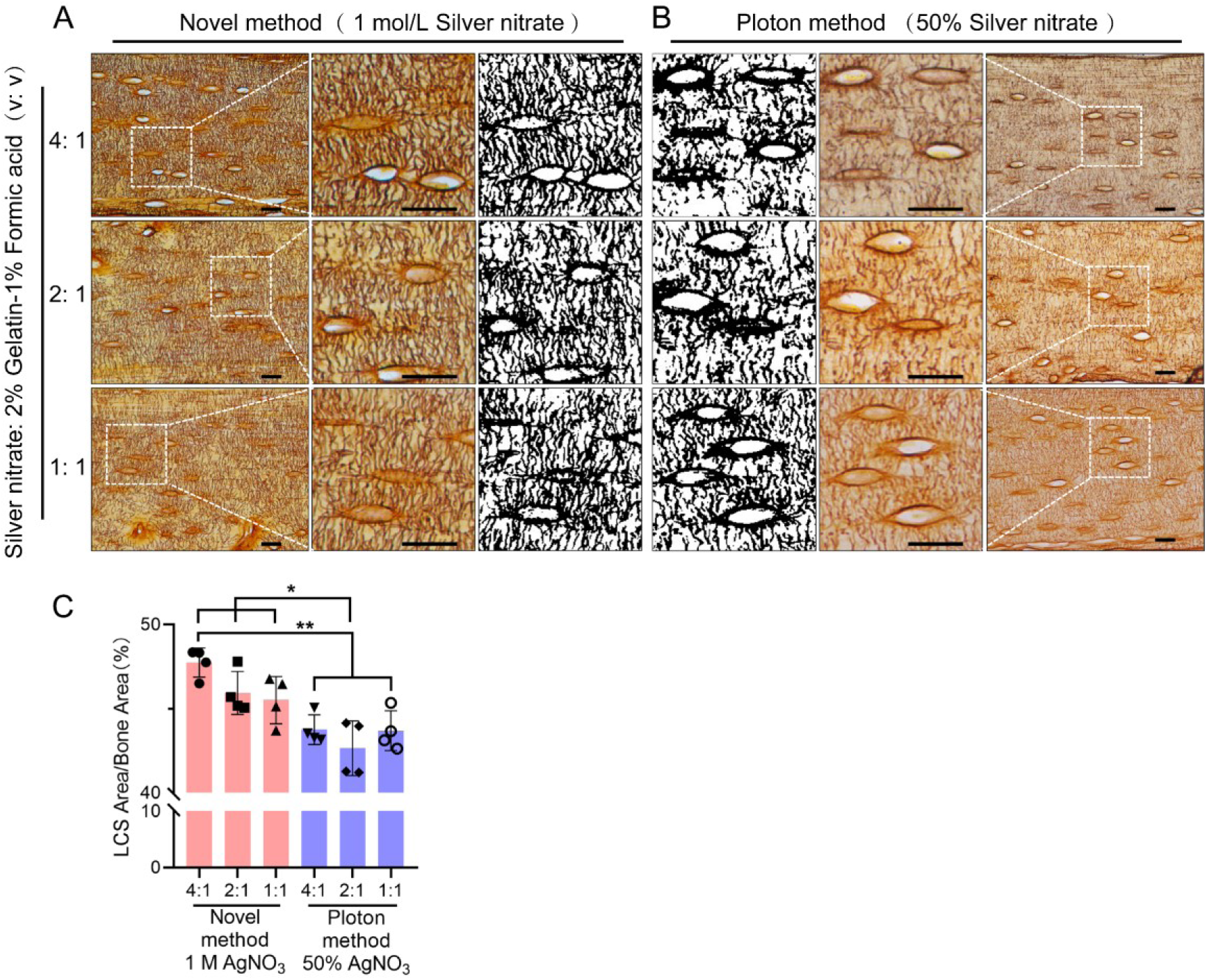
Comparison of the Staining effects of 1 mol/L Silver Nitrate and 50% Silver Nitrate on LCS. (A) Staining using 1 mol/L Silver Nitrate and 2% gelatin-1% formic acid at different volume ratios (4:1, 2:1, 1:1), with the third column showing binarized images. Scale bars = 20 μm. (B) Staining using 50% Silver Nitrate and 2% gelatin-1% formic acid at different volume ratios (4:1, 2:1, 1:1), with the first column showing binarized images. Scale bars = 20 μm. (C) Comparison of staining effects on osteocyte LCS density between the Novel method and the Ploton method, analyzed using one-way ANOVA. Data are presented as mean ± SD. *p < 0.05; **p < 0.01, n = 4 for repeated staining with 3-4 ROIs per staining section.

**Figure 7.**
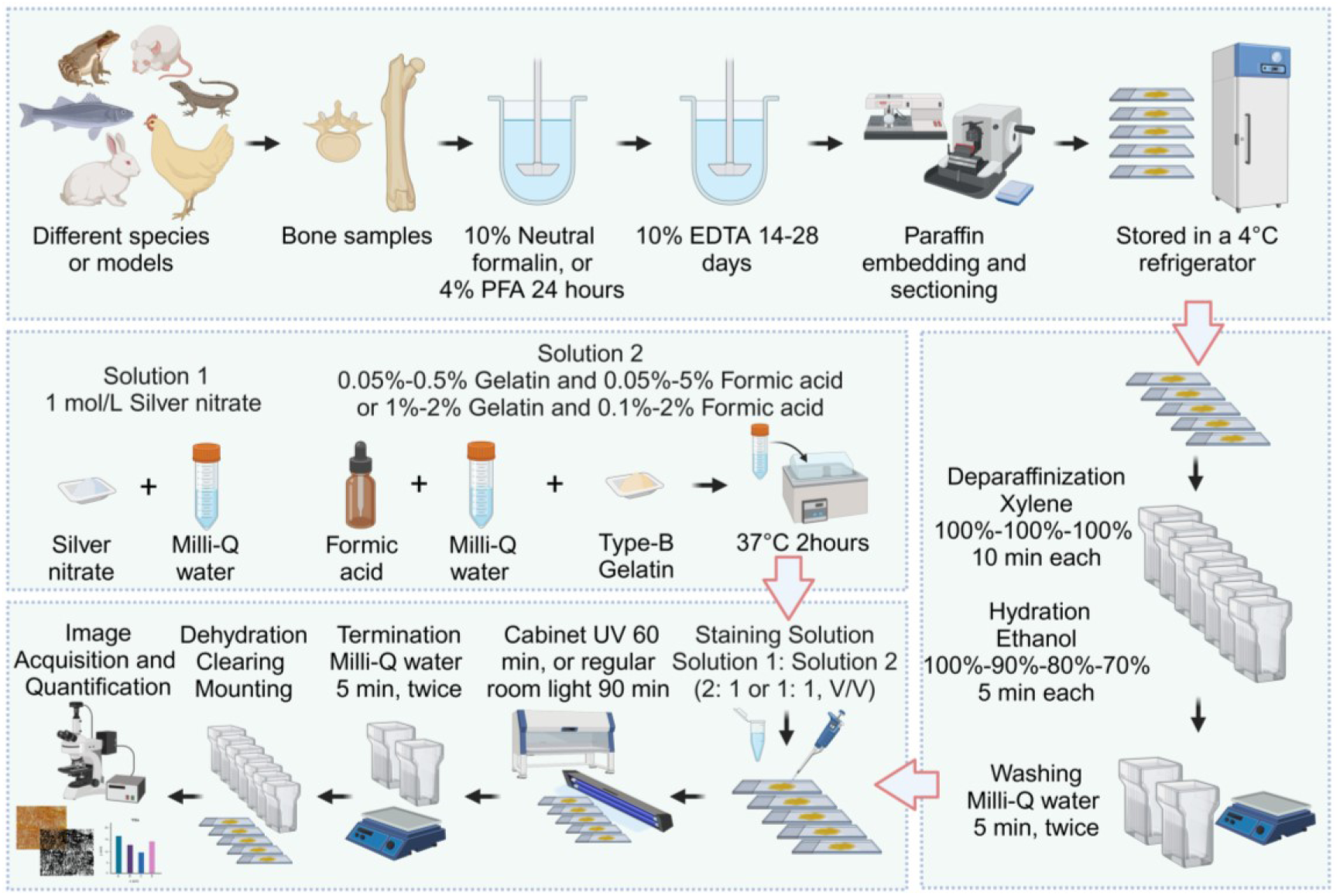
Flowchart of the Novel Optimized Silver Nitrate Staining Method. This schematic outlines the key steps for staining the osteocyte LCS. Created in BioRender. Libo, W. (2024) https://BioRender.com/m59m214

The results showed that the density of LCS stained with 1 mol/L silver nitrate was significantly higher than that stained with 50% silver nitrate, particularly when the volume ratio of silver nitrate solution to 2% gelatin-1% formic acid solution was 4:1 or 2:1. The LCS stained with 1 mol/L silver nitrate appeared more intact and clear (Figure 6A), whereas the LCS stained with 50% silver nitrate was more intermittent, irregular, and exhibited blurred contrast(Figure 6B).

In addition, statistical analysis shows that when using the classic Ploton staining solution ratio (i.e., a 2:1 volume ratio of 50% silver nitrate to 2% gelatin-1% formic acid solution), the LCS density stained is significantly lower than that stained with a 1 mol/L silver nitrate and 2% gelatin-1% formic acid solution at volume ratios of 4:1, 2:1, and 1:1(Figure 6C). This result suggests that the higher concentration of silver nitrate may cause damage to the LCS during the staining process, or that the rapid reaction rate induced by the high concentration does not allow for complete staining of all LCSs. We hypothesize that these factors contribute to the superior staining effectiveness of 1 mol/L silver nitrate compared to the Ploton method, which uses 50% silver nitrate.

## Discussion

In this study, we developed and optimized a novel silver nitrate staining method for characterizing and quantifying the osteocyte LCS. Our results demonstrate that this optimized method, utilizing 1 mol/L silver nitrate, stains and quantifies the osteocyte LCS more efficiently than the classical Ploton method, which uses 50% silver nitrate.

Impregnation staining with a high concentration of silver nitrate (50%, w/v) is the most widely used technique for characterizing and quantifying the osteocyte LCS. This method was first reported by Goodpasture and Bloom in 1975 for staining chromosomal nucleolus organizer regions (NORs) [24], adapted from Howell et al.’s study published the same year, which focused on staining the satellite regions of human chromosomes with a 10% silver nitrate solution [25]. In 1980, Howell and Black simplified Goodpasture and Bloom’s three-step staining method into a single step, improving the reproducibility and operability of the technique [15]. Notably, the solutions used in their one-step method—50% silver nitrate in deionized water (Solution 1) and 2% gelatin in 1% formic acid (Solution 2), in a 2:1 volume ratio—remain unchanged today, and the incorporation of gelatin, a protective colloid, helped control the silver staining process and reduce background staining caused by the intense reduction of silver nitrate. In their 1986 study, Ploton et al. lowered the reaction temperature from 70°C to 20°C and used a 5% sodium thiosulfate solution to terminate the staining process [16], whereas Howell and Black’s method only used deionized water for washing out the staining mixture. In these studies, the staining duration was brief, ranging from a few seconds to a few minutes, as the staining was conducted on chromosomes in metaphase during mitosis. According to the available literature, Chappard et al. first applied Ploton method in 1996 to stain the nucleolus organizer regions (NORs) of cells in undecalcified, MMA-embedded bones [17, 26]. During this process, they accidentally discovered that the method also provided excellent staining of the osteocyte LCS. It was Chappard et al. who first determined that the optimal staining time for the osteocyte LCS was "55 minutes at room temperature in the dark," followed by washing the slides with a 5% sodium thiosulfate solution. These conditions—both the staining reaction and the termination procedure—have since become the standard protocol for staining the osteocyte LCS. In subsequent studies, Chappard et al. adapted this method in staining the osteocyte LCS in paraffin-embedded bone samples after decalcification [27–28].

Over the last two decades, there has been a growing recognition of the essential role osteocytes play in bone remodeling, the remodeling of the lacunar-canalicular system, and the development of osteocyte dendrites [29–32]. Researchers have also adopted the staining solutions first described by Howell and Black, along with the 5% sodium thiosulfate solution for termination and room temperature reaction conditions introduced by Ploton et al., as well as the reaction time and dark conditions outlined by Chappard et al. This method is widely recognized in osteocyte research as Ploton silver staining [18] or the Ploton impregnation method [28].

In our preliminary study, we encountered difficulties in obtaining sufficient silver nitrate powder to prepare a 50% silver nitrate solution for osteocyte biology studies due to safety regulations. Instead, we utilized a 1 mol/L silver nitrate solution, which is readily available for chemical titrations, as a substitute for the 50% solution in staining the osteocyte LCS. Surprisingly, this solution effectively and clearly stained the osteocyte LCS. This finding prompted us to optimize the classical Ploton method by using a lower concentration of silver nitrate solution, thereby enabling more efficient and environmentally friendly staining and quantification of the osteocyte LCS.

Based on this idea and using the classical Ploton method as a reference, we evaluated several factors that could influence the effectiveness of silver nitrate staining. Consequently, we selected six key conditions for study: silver nitrate concentration, the concentrations of the gelatin-formic acid combination, the volume ratio of silver nitrate to gelatin-formic acid solution, type of gelatin, light conditions, and termination conditions. Our results demonstrated that, for 4 μm paraffin bone sections, optimal staining was achieved with a 1 mol/L silver nitrate solution combined with a gelatin-formic acid solution at various concentrations (0.05%-0.5% type-B gelatin and 0.05%-5% formic acid, or 1%-2% type-B gelatin and 0.1%-2% formic acid) in volume ratios of 4:1, 2:1, or 1:1, or a 0.5 mol/L silver nitrate solution at a 4:1 ratio. The staining process was carried out for 1 hour under UV light in a biosafety cabinet or 90 minutes under regular room light (or dark), followed by washing with Milli-Q water to terminate the reaction.

Next, we applied the optimized, low-concentration, cost-effective, and environmentally friendly staining method to the osteocyte LCS in bone samples from various animal species. Our experiments demonstrated that this method effectively provides clear contrast staining of the osteocyte LCS in the bones of mammals, amphibians, fish, birds, and reptiles. Notably, we observed that teleost vertebrae, compared to those of other species, contained significantly fewer mature osteocytes and exhibited a notably lower number and density of LCS. This finding suggests that the optimized staining method could serve as a powerful tool for comparative evolutionary studies of osteocyte and LCS morphogenesis across different vertebrate taxa.

In addition, the optimized method effectively stains the LCS in both glucocorticoid-induced bone growth retardation mice and Zeb1 overexpression transgenic mice (Prrx1Cre;Zeb1*^fx/fx^*). In the model group, the density of the LCS in the femoral cortical bone was significantly lower than in the control group. Notably, in the Prrx1Cre;Zeb1*^fx/fx^* mice, we observed not only a marked reduction in LCS density but also extensive cartilage remnants in the cortical bone (Alcian Blue-Safranin O staining, data not shown). These findings, for the first time, suggest that the Prrx1Cre;Zeb1*^fx/fx^*transgenic mice may serve as a new model for osteogenesis imperfecta, providing a valuable platform for investigating the expression patterns and underlying mechanisms of transcription factors involved in bone development and osteocyte maturation.

Finally, we compared the optimized method (1 mol/L silver nitrate) with the classic Ploton method (50% silver nitrate) for LCS staining. Statistical analysis revealed that the 1 mol/L silver nitrate staining resulted in significantly higher LCS density compared to the 50% silver nitrate method. This finding highlights a potential limitation of the Ploton method: the 50% silver nitrate solution may not fully reveal osteocyte LCS in certain bones. The strong reduction reaction of the high-concentration silver nitrate could damage the LCS or fail to adequately stain osteocyte LCS in certain species or models, potentially leading to an underestimation of osteocyte LCS density or number. Although no direct comparison was made in this study, we also observed that the optimized method yielded better results than another technique using the Silver-Protein method (or Bodian’s Protargol-S method) for staining osteocyte LCS [33, 34]. We recommend that researchers studying osteocytes carefully consider the inherent limitations of the Ploton method and Silver-Protein method, and critically evaluate the novel optimized method presented in this paper.

## Conclusion

In conclusion, this study represents the first systematic optimization of a staining method that has been established for nearly half a century and utilized in osteocyte research for almost 30 years. The novel optimized method uses a 1 mol/L silver nitrate solution combined with a gelatin-formic acid solution at various concentrations (0.05%-0.5% type-B gelatin and 0.05%-5% formic acid, or 1%-2% type-B gelatin and 0.1%-2% formic acid) in volume ratios of 4:1, 2:1, or 1:1, or a 0.5 mol/L silver nitrate solution at a 4:1 ratio. The staining process was carried out for 1 hour under UV light or 90 minutes under regular room light (or dark), followed by washing with Milli-Q water to terminate the reaction. This novel method effectively stains LCS across different species and disease models, demonstrating greater number and density of stained LCS compared to the classical Ploton method. We believe that our optimized method will advance osteocyte biology research and provide a valuable tool for studying the adaptive evolution of osteocyte LCS morphology and function.

## Acknowledgments

This research was funded by grants from the National Natural Science Foundation of China (NSFC) (82104574, 82405480, 82374474), 5^th^ Long-Yi Scholar Project of Longhua Hospital Shanghai University of Traditional Chinese Medicine (PY2022001), Long-Yi S&T Innovation Cultivation Project of Longhua Hospital Shanghai University of Traditional Chinese Medicine (YD202208), Young Talent Training System of Longhua Hospital Shanghai University of Traditional Chinese Medicine (SZLRZ-2024-38), Traditional Chinese Medicine Science and Technology Development Project of Shanghai Medical Innovation & Development Foundation (No.WL-HBMS-2022003K).

## CRediT authorship contribution statement

Wu Jinlian and Xue Chunchun: Data curation, Investigation, Validation, Funding acquisition, Writing-original draft. Li Qiang and Wang Chenlong: Resources. Zhang Jie and Wu Hongjin: Formal analysis. Dai Weiwei and Wang Libo: Conceptualization, Methodology, Supervision, Funding acquisition, Project administration, Writing-review & editing.

## Dedication

Wu Jinlian and Wang Libo dedicate their first scientific discovery to their beloved parents and daughters, Luna and Lina, in gratitude for their cherished love and unwavering support throughout their academic pursuit.

